# Shaping nanoparticle diffusion through biological barriers to drug delivery

**DOI:** 10.1101/2021.07.21.453209

**Authors:** Benjamin J. Lee, Yahya Cheema, Shahed Bader, Gregg A. Duncan

**Author notes:** Correspondence to: Gregg Duncan.

## Abstract

Nanoparticle drug delivery systems encounter many biological barriers, such as the extracellular matrix and mucus gels, that they must bypass to gain access to target cells. A design parameter that has recently gained attention is nanoparticle shape, as it has been shown elongated rod–shaped nanoparticles achieve higher diffusion rates through biological gels. However, the optimal dimensions of rod-shaped nanoparticles to enhance this effect has yet to be established. To systematically approach this, rod-shaped nanoparticles were synthesized by mechanically stretching 100 nm, 200 nm, and 500 nm spherical nanoparticles. Transmission electron microscopy confirmed this procedure yields a significant fraction of elongated rods and remaining spheres could be removed by centrifugation. Fluorescent microscopy and multiple particle tracking analysis was then used to characterize rod-shaped and spherical nanoparticle diffusion in MaxGel®, a model extracellular matrix hydrogel. When dispersed in MaxGel, we found rod-shaped nanoparticles exhibited the greatest enhancement in diffusion rate when their length far exceeds the average hydrogel network size. These results further establish the importance of shape as a design criterion to improve nanoparticle-based drug delivery systems.

## Introduction

Nanomedicine provides a promising alternative to traditional therapies for efficient drug delivery to selectively target tissues and as contrast agents for diagnostic imaging applications.^1^ Furthermore, nanomedicine-based pharmaceuticals have demonstrated reduced immunogenic and off-target side effects common with traditional therapies.^1, 2^ For pulmonary drug delivery applications, the inhaled administration route has been a target of interest to deliver nanotherapeutics given the high surface area and extensive vascularization of the lung.^4–6^ Despite these benefits, there are various obstacles to be overcome, which if not accounted for can hinder the clinical performance of nanomedicine.^2^ In order to achieve the desired therapeutic outcomes, nanomedicine must efficiently diffuse through and avoid adhesion to biological gels, such as the extracellular matrix (ECM) and mucus, in order to reach target cells within the tissue of interest. For example, ECM possesses a mesh-like structure through the cross-linking and entanglement of collagen fibers and other proteoglycans which can inhibit efficient transport and reduce retention of nanoparticle drug delivery systems.^3–5^ In diseased states such as cancer, the ECM becomes more concentrated making this microenvironment less easily penetrated and presenting a greater barrier to drug delivery.^6^ Likewise, the biodistribution and efficacy of inhaled drug and gene nanocarriers can be significantly limited by mucus secreted on the airway epithelial surface depending on nanoparticle size and surface chemistry.^7, 8^ In addition, mucus observed in diseases such as cystic fibrosis and asthma is significantly more concentrated and densely cross-linked, which may further inhibit efficient pulmonary nanoparticle drug delivery.^9, 10^ Taken together, extracellular barriers within target tissues are likely to influence the effectiveness of nanoparticle-based therapies.

Many pathogenic viruses and bacteria have complex, anisotropic shapes which enhance their ability to survive and replicate in their host.^11, 12^ These observations have inspired research exploring shape as a design parameter for drug delivery systems.^13, 14^ Prior work has shown rod-shaped nanoparticles can more readily penetrate through biological barriers to delivery. For example, it has been shown rod-shaped tobacco mosaic viral filaments achieves more rapid diffusion through 3D tumor spheroids.^15^ In addition, previous studies have shown that mesoporous silica nanorods exhibit higher diffusion in gastrointestinal (GI) mucus and longer retention in the GI tracts of mice than its spherical counterparts when orally administered.^16, 17^ Using molecular simulations, they found that nanorods better penetrate GI mucus due to non-uniform rotational motion and directional transport under shear flow.^16^ For targeted drug delivery applications, it has been demonstrated ligand-functionalized, rod-shaped nanoparticles possess highly specific adhesion to lung and brain endothelium in an *in vitro* microfluidic model mimicking the vasculature and in an *in vivo* model where mice were injected with intercellular adhesion molecule monoclonal antibody (ICAM-mAb)-coated nanorods.^18, 19^

While these important findings support the use of rod-shaped nanomaterials for biomedical applications, prior studies have yet to systemically characterize rod-shaped nanoparticle behavior in biological barriers to drug delivery, such as the ECM. Herein, we report measurements of nanorod and nanosphere of varying size to find the optimal dimensions for nanoparticle penetration through a model ECM hydrogel. Elongated rod-shaped nanoparticles were prepared using a previously established mechanical stretching technique.^20^ The motions of individual nanoparticles were directly measured using fluorescent microscopy and multiple particle tracking analysis. We show here that nanorod dimensions must be tuned with respect to ECM architecture where increasing nanorod length beyond the ECM mesh pore size leads to marked enhancements in diffusion rate. The diffusion of nanorods with lengths less than or in range of the ECM mesh spacing was left unchanged or reduced as compared to their spherical counterparts. These data provide new insights on the shape dependence of nanoparticle dynamics in biological gels and design considerations in creating nanoparticle drug delivery systems.

## Methods

### Mechanical stretching of nanospheres (NS) into nanorods (NR)

The protocol described here was adapted from a previously reported method.20 To create a film with embedded NS, 20 mL of 5% w/v solution of polyvinyl alcohol (PVA) was mixed using a magnetic stir bar and heated at 130°C for at least one hour until fully dissolved. The solution was passed through a fine mesh grid to remove any undissolved lumps of PVA. Next, 100 nm, 200 nm, or 500 nm fluorescent carboxyl-coated polystyrene NS (ThermoFisher) were mixed into the PVA solution at concentration ranging from 0.01–0.1% w/v. To plasticize the film, 320 μL of glycerol was also added to the PVA/NS containing solution. The solution was then poured into a 12 × 12 cm mold and left to dry overnight uncovered. Small bubbles present in the film were removed using tweezers. Once the film fully dried, two 9 × 5 cm films were cut from the film and submerged in toluene overnight to liquefy the nanoparticles. Films were taken out, and a 5 × 5 cm grid was drawn on the center of the film using a marker to visually determine the uniformity of the stretching. The films were loaded on an Arduino-controlled mechanical stretcher by adhering and clamping the film on the platform using tape and Gorilla Glue. The film was then stretched to up to 3 times the size of the marked grid and left to dry overnight. The film was cut along the grid marks and any non-uniformly stretched portions were discarded. The remaining film sections were submerged in isopropanol to remove any residual toluene. After 24 hours, the film sections were removed and then put into a 30% isopropanol solution, which was then mixed and heated at 130 °C for an hour until the film had fully dissolved. The dissolved films were centrifuged for 30 minutes at 26,000xg, and the supernatant was discarded. The pellet was re–suspended in 30% isopropanol at 130 °C and after 1 hour, centrifuged at 26,000xg. This process of PVA film dissolution and washing was repeated three times. To separate the rods from the spheres, solutions of stretched NS and NR were put into 1.5 mL centrifuged tubes and spun at 12,000xg for 100 nm and 200 nm NS and 10,000xg for 500 nm NS for 10 minutes. The pellet that accumulated on the side, theoretically containing NR, was disrupted using a spatula and the supernatant was decanted into a separate centrifuge tube.^21^ The remaining pellet containing NS at the bottom of the tube was resuspended in 500μL upH_2,_O.

### Transmission Electron Microscopy (TEM)

Stretched nanoparticles were diluted 100x in ultrapure H_2,_O and 25 μL was placed on a square of Parafilm. A PELCO® copper mesh grid (400 mesh Cu, Ted Pella) was inverted and placed on this droplet for 30 seconds. After, the grid was washed 4 times for 30 seconds by placing on a 25 μL droplet of ultrapure H_2_O. After drying, the grid was imaged using transmission electron microscopy (JEM-2100, 200 kV, JEOL Ltd).

### Fluorescent video microscopy and multiple particle tracking (MPT) analysis

Custom microscopy chambers were prepared as previously described.^22^ Microscopy chambers were either filled with H20 for control experiments or ~1 mg/mL MaxGel (Sigma Aldrich). Slides containing MaxGel were incubated at 37°C for 30 minutes to form a gel. After incubation, 1 μL of ~1000x diluted NS or NR was added to 24 μL of sample. The slides were covered with a plastic cover slide and left for 30 minutes to allow the nanoparticles to equilibrate in the solution. Slides containing MaxGel were left in the incubator until ready for MPT analysis. Fluorescence video microscopy and MPT analysis was used to determine the translational diffusion of the nanoparticles in aqueous and hydrogel media. MPT experiments were performed using a Zeiss 800 LSM microscope and videos were acquired at 33.33 Hz frame rate for 10 seconds. Videos were analyzed using TrackPy (*github.com/soft-matter/trackpy*), a Python-based particle tracking code that detects Gaussian blob-like features in a video. Particle trajectories are then used to calculate time-averaged mean square displacement (MSD) as a function of lag time, τ and from this, the average translational diffusion coefficient (*D*) of the nanoparticles can be determined. For a nanoparticle in a Newtonian fluid, *D* for NS can be calculated from the Stokes-Einstein equation,

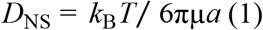

where *k*_B_ is Boltzmann’s constant, *T* is the temperature, μ is the viscosity of the solution, and *a* is the radius of the nanoparticle. When calculating the theoretical rate of diffusion for rod-shaped nanoparticles in a purely viscous fluid, a modified version of the Stokes-Einstein Equation must be used to account for anisotropic hydrodynamic effects. Specifically, the diffusion coefficient for NR (*D_NR_*) must account for motion parallel and perpendicular with respect to the long axis and can be calculated as,

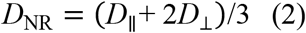

where *D*_∥_ = (*k*_B_*T*/2πμ) × ln[*L/d*], *D*_⊥_ = (*k*_B_*T*/4πμ) × ln[*L/d*], *L* is the length of the long axis of the nanorod, and *d* is the cross-sectional diameter of the shorter axis.^23^ Additional analyses were performed to characterize the viscoelastic properties MaxGel through MPT. The logarithmic slope of MSD (α) can be calculated as α =log_10_(⟨MSD(τ)⟩)/log_10_(τ). For a gel network, sub-diffusive nanoparticle movement should be observed where 0 < α <1.^24^ Using the generalized Stokes-Einstein relation, measured MSD values were used to compute viscoelastic properties. The Laplace transform of ⟨MSD(τ)⟩, ⟨MSD(s)⟩, is related to viscoelastic spectrum 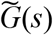 using the equation 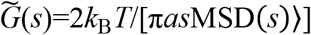, where *s* is the complex Laplace frequency. The complex modulus can be calculated as *G*(ω)* =*G*′(*ω*) +*G*′′(*iω*), with iω being substituted for *s,* where *i* is a complex number and is frequency.^24^ Hydrogel network pore size, ξ, is estimated based on *G*’ using the equation, *ξ*≈(*k*_B_*T*/G’)^1/3^. ^24^

## Results & Discussion

### Mechanical Stretching of Nanoparticles into Ellipsoidal Rod-Shapes

In this work, rod-shaped ellipsoidal nanoparticles were fabricated by unidirectional mechanical stretching of spherical nanoparticles. Films containing fluorescent, carboxyl-coated polystyrene NS of diameters 100 nm, 200 nm, and 500 nm were stretched by ~3X their original length. TEM was used to determine the overall uniformity and shape of the nanoparticles after stretching (**Fig. 1**). TEM micrographs show that the procedure yielded a significant number of rod-shaped ellipsoidal particles, termed here as “nanorods” (NR). The length and width of NR were determined using ImageJ (**Table 1**).

**Table 1.** Average length (*L*) and width (*d*) of 100nm, 200nm, and 500nm nanoparticles after stretching The number of nanoparticles (n) measured and standard deviation in nm are also indicated.

|  | Original Nanoparticle Diameter |  |  |
| --- | --- | --- | --- |
|  | 100 nm (n = 109) | 200 nm (n = 105) | 500 nm (n = 99) |
| $L$ (nm) | $302 \pm 18.4$ | $705 \pm 132$ | $1770 \pm 208$ |
| $d$ (nm) | $58 \pm 7$ | $126 \pm 21$ | $285 \pm 16$ |

**Figure 1.**
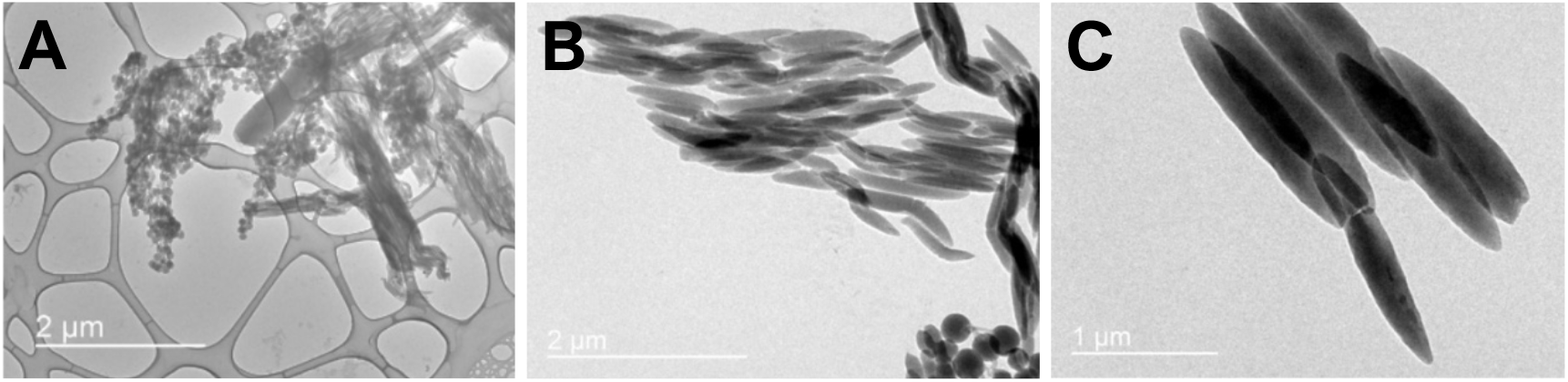
Ellipsoidal nanorod production. Transmission electron microscopy images of (**A**) 100 nm, (**B**) 200 nm, and (**C**) 500 nm polystyrene nanospheres after mechanical stretching into ellipsoidal nanorods.

### Centrifugal separation of nanorods and nanospheres

Based on TEM micrographs (**Fig. 1**), it was apparent a significant fraction of spherical nanoparticles remained after stretching. In order to generate more purified NR stocks, we adapted a procedure previously shown to separate gold NS and NR by a simple centrifugation step.^21^ Specifically, stretched samples containing both NR and NS were centrifuged to create shape-dependent pellets where NS accumulate on the bottom and NR accumulate on the sides of the centrifuge tube (**Fig. 2A**).^21^ After the pellets were formed, NR on the side were carefully removed from the wall and put into a separate tube. NS within the bottom pellet were re-dispersed in water in the same tube. It was also apparent the amount of unstretched particles remaining was most significant after stretching of 100 nm NS (**Fig 1A**). As such, we subjected these 100 nm–based NR samples to MPT analysis to assess the uniformity of the recovered NS fraction based on their diffusion in water as compared to stock NS. MSD of the NS recovered by centrifugation and stock NS were calculated and diffusion coefficients were determined based on linear regression (**Fig. 2B**). We find the MSD and diffusion coefficient are very similar for stock and recovered NS with a predicted diameter of 115 and 114 nm, respectively (**Eq. 1**). We next compared the MSD and diffusion rate of NR recovered by centrifugation to stock NS (**Fig. 2C**). We find, as expected, NR diffuse more slowly than NS in water based on measured MSD. Assuming an aspect ratio (*L*/*d*) of 5, we fit for an effective diffusion coefficient for the NR using **Eq. 2** and find the predicted dimensions of NR (L=355 nm; d=71 nm) are similar in magnitude to our TEM measurements (**Table 1)**. These results support centrifugal separation of stretched NS sample yields a higher purity of NR than what was observed through TEM. As such, we used centrifugal purification in our subsequent studies on NR diffusion in ECM hydrogels.

**Figure 2.**
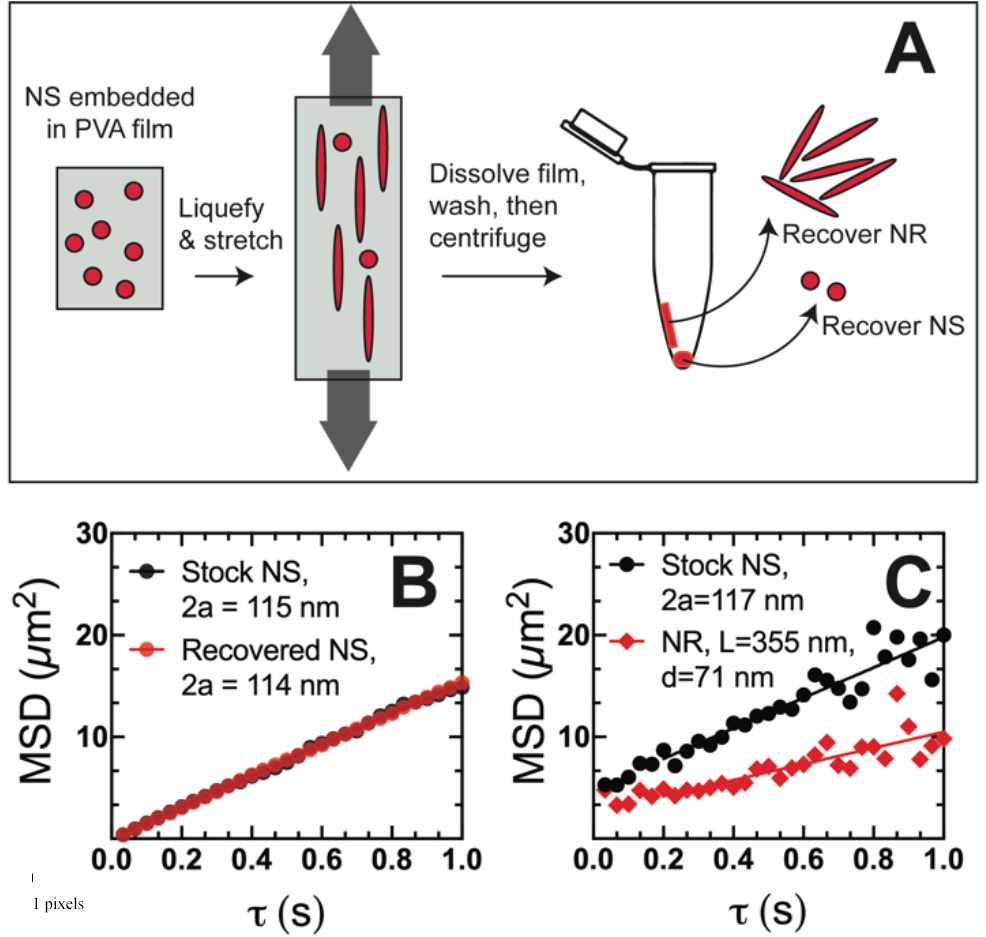
Diffusion of stretched nanorods (NR) and unstretched nanospheres (NS) isolated by centrifugation compared to as-prepared stock NS. **(A)** Illustration of the stretching procedure and isolation of nanospheres (NS) and nanorods (NR) by centrifugation. (**B, C**) Multiple particle tracking was conducted for NS and NR in water to determine their respective sizes based on measured mean squared displacement (MSD) and diffusion rate (*D*). (**B**) MSD versus time lag (τ) of unstretched stock NS and recovered spherical nanoparticles (NS) separated by centrifugation in water. (**C**) MSD versus time lag (τ) of unstretched stock NS and recovered NR separated by centrifugation. The diameter (2*a*) of NS was calculated from measured *D*_NS_ using Equation 1. The length (*L*) and diameter (*d*) of NR were determined based on measured *D*_NR_ using Equation 2 assuming an aspect ratio (*L*/*d*) of 5.

### Microrheology and network structure of ECM hydrogels (MaxGel)

To gain insights on the behavior of NR and NS in an ECM-like environment, we chose to use MaxGel as a model as it is composed of fibroblast–derived, human ECM proteins such as collagen, laminin, and other glycosaminoglycans. MaxGel has been used in prior studies as a tissue culture scaffold to recapitulate the ECM microenvironment.^25, 26^ In order to characterize the microscale mechanical properties of MaxGel, we employed MPT microrheology using 100 nm nanoparticles coated with polyethylene glycol (PEG), prepared as described in previous work^22^ (**Fig. 3**). We find based on measured MSD that these nanoparticles exhibited sub-diffusive motion based on the logarithmic slope of MSD versus time, α=log[MSD]/log[τ] < 1 (**Fig. 3A**). Sub-diffusive motion is expected for nanoparticles embedded within a 3D polymer network. Next, we quantified the microrheological properties of MaxGel using the generalized Stokes-Einstein equation to find microscale elastic (G’) and viscous (G”) moduli (**Fig. 3B,C**). Both G’ and G” increased as a function of frequency and G’ exceeds G” at all measured frequencies which is characteristic of hydrogel networks with viscoelastic properties. However, we should note our measurements are limited to relatively low frequencies (*ω* ≤ 1 Hz) using MPT microrheology.^24^ We estimated network size (*ξ*) within MaxGel based on measured G’ at *ω* = 1 Hz for individual nanoparticles (**Fig. 3D**) which varied over a range between ~200–600 nm (*ξ*_avg_ = 415 nm). This heterogeneity in measured *ξ* is expected for a multi-component ECM hydrogel. These measurements are important as they provide a basis for comparison between the NR and NS of varying size to MaxGel network dimensions.

**Figure 3.**
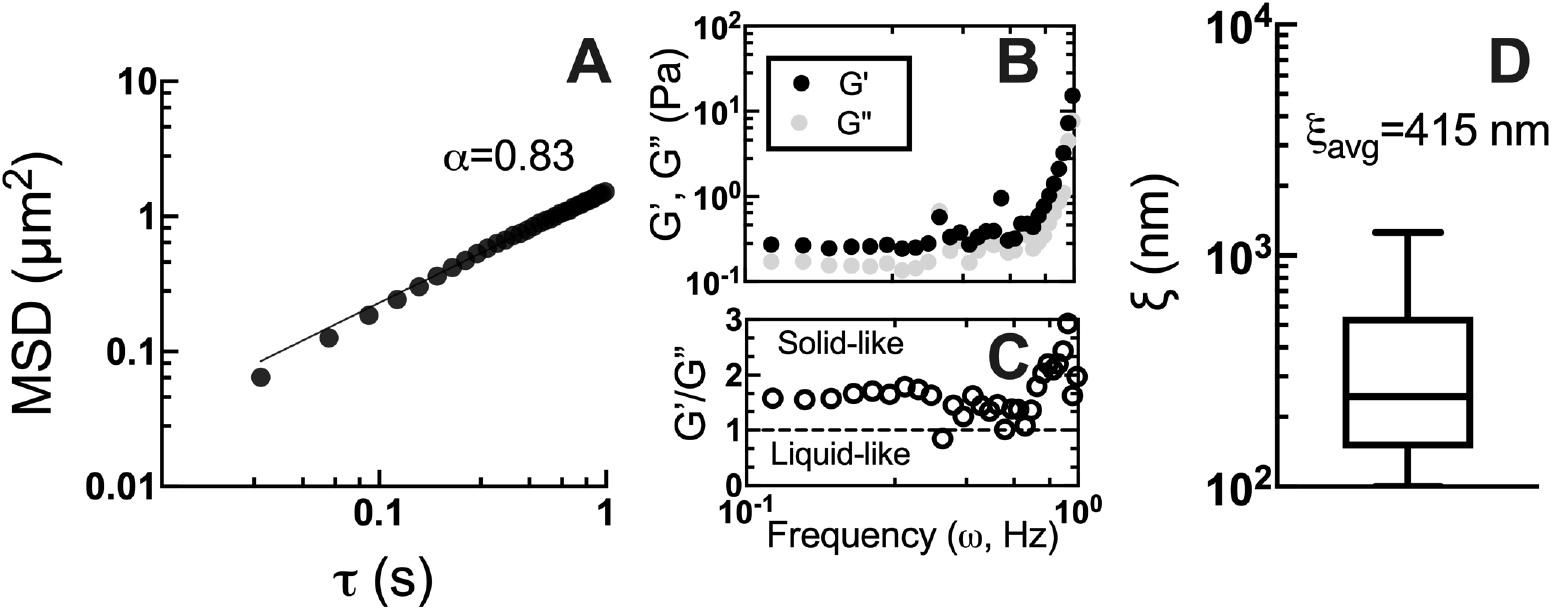
Microscale viscoelastic properties of ECM hydrogels (MaxGel). **(A)** Mean squared displacement (MSD) versus time with log-log scaling for PEGylated 100 nm NS in MaxGel. The logarithmic slope of the MSD (α=log[MSD]/log[τ]) is also indicated where values below 1 are indicative of sub-diffusive motion. (**B**) Elastic (G’) and viscous moduli (G”) and (**C**) G’/G” as a function of frequency (ω) based on microrheological analysis of measured MSD. Values of G’/G” greater than 1 are indicative of elastic solid-like behavior and values less than 1 are indicative of viscous liquid like behavior. (**D**) Estimated pore size (*ξ*) based on analysis of *G*′ at *ω* = 1 Hz calculated as *ξ* ≈ (*k*_B_*T*/*G*′)^1/3^.

### Shape-dependent nanoparticle diffusion in ECM hydrogels (MaxGel)

Finally, the diffusion of NR generated by mechanical stretching and as-prepared stock NS were measured in MaxGel using MPT. We note NS and NR were added to solidified MaxGel following 30 minutes of incubation at 37°C. The NS and NR were dispersed within the gel by mechanical mixing and were allowed to equilibrate for an additional 30 minutes at 37°C prior to imaging. In addition, separate micro-well chambers containing MaxGel were prepared for each NS and NR tested as they fluoresce at the same wavelength. As we observed non-uniform behavior of NS and NR within MaxGel, we compared the overall distribution of MSD at τ =1 s (MSD_1s_) for NS and NR (**Fig. 4**). Given their comparable surface areas, we compared the diffusion of stock NS to NR prepared using NS of the same size. Interestingly, we find 100 nm NS and stretched NR (L≈300 nm, d≈60 nm) showed comparable diffusion rates (**Fig. 4A**). Further, 200 nm NS-based NR (L≈700 nm, d≈125 nm) were more restricted within MaxGel as compared to the original 200 nm NS (**Fig. 4B**). Conversely, we observed an enhancement in diffusion rate for the largest 500 nm NS-based NR (L≈1.8 μm, d≈285 nm) as compared to 500 nm NS (**Fig. 4C**). For the NS, we observed the expected trend that smaller NS generally diffused faster than larger NS. However, when comparing NR, the ~1.8 μm long NR diffused the most rapidly overall and ~700 nm long NR exhibited the slowest diffusion rate.

**Figure 4.**
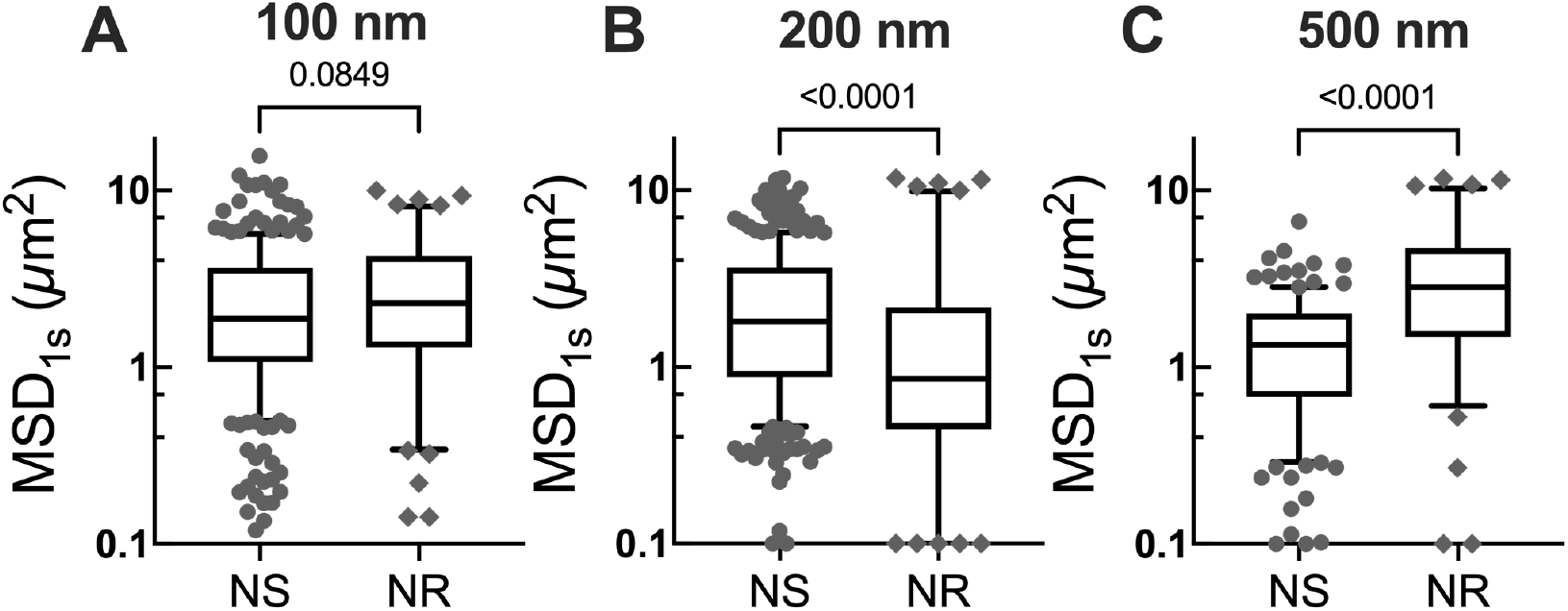
Nanosphere and nanorod diffusion within ECM hydrogels (MaxGel). Measured mean squared displacement at τ =1 s (MSD_1s_) of nanospheres (NS) and nanorods (NR) within MaxGel. Data are shown for NS and NR with original diameters of (**A**) 100 nm, (**B**) 200 nm, and (**C**) 500 nm in MaxGel. Experiments were repeated at least in triplicate with 10–20 videos analyzed and 100–300 individual particles tracked per condition. Mann-Whitney U test was used to compare NS and NR MSD_1s_ where *p*– value less than 0.05 was considered significant.

These results can be interpreted based on prior experimental and computational studies demonstrating constrained motion perpendicular to the long axis can lead to enhancements in translational diffusion through a polymer mesh network.^24, 27, 28^ For ~300 nm NR, their motion is comparable to 100 nm NS given their overall dimensions are smaller than the average network spacing in MaxGel (d < L < ξ_avg_). For ~700 nm NR, although in the size range (d < ξ_avg_ < L) where we would predict faster diffusion due to reduced frictional drag parallel to the long axis, we do not observe enhanced diffusion which may be due in part to the natural heterogeneity of the ECM network and wide range of pore size which may interfere with these effects. The larger ~1.8 μm NR are in the predicted regime where NR diffusion is increased due to reduced hydrodynamic drag parallel to the long axis based on the comparative size of NR and ξ (d < ξ_avg_ ≪ L). These observations would suggest that NR length should far exceed network pore sizes within the ECM to achieve enhanced penetration through the ECM. However, additional work is required to determine how disease-associated changes to ECM (e.g. increased concentration and/or cross-linking) would impact NR penetration and if these results are generalizable to other biological barriers (e.g. mucus). It will also be of interest in future work to determine if these findings may apply to flexible NR^29, 30^ as compared to the rigid NR studied here and how surface chemistry of NR influences diffusion behavior within ECM and other biological barriers to drug delivery.^31, 32^

## Conclusion

This work provides new insights into how shape influences the dynamic behavior of nanoparticles within a model ECM hydrogel. While our stretching procedure yielded a mixture of rods and spheres, we demonstrated rod-shaped particles could be purified using centrifugation. Clear differences were observed in the diffusion of rod-shaped and spherical nanoparticles within MaxGel which were dependent on how the length of rod-shaped nanoparticles compared to the ECM mesh spacing. Our results underscore the significance of the ECM as a barrier to nanoparticle drug delivery systems and the potential benefit of controlling shape to improve their distribution within target tissues.

## Acknowledgements

This work was supported by the University of Maryland, the Burroughs Wellcome Fund Career Award at the Scientific Interface, the American Lung Association Innovation Award, and the NSF CAREER Award 2047794. We also acknowledge the support of the Maryland NanoCenter and its AIMLab for access to and assistance with TEM imaging.

